# Automated liquid-handling operations for robust, resilient, and efficient bio-based laboratory practices

**DOI:** 10.1101/2022.11.03.515004

**Authors:** Mario A. Torres-Acosta, Gary J. Lye, Duygu Dikicioglu

## Abstract

Increase in the adoption of liquid handling devices (LHD) can facilitate experimental activities. Initially adopted by businesses and industry-based laboratories, the practice has also moved to academic environments, where a wide range of non-standard/non-typical experiments can be performed. Current protocols or laboratory analyses require researchers to transfer liquids for the purpose of dilution, mixing, or inoculation, among other operations. LHD can render laboratories more efficient by performing more experiments per unit of time, by making operations robust and resilient against external factors and unforeseen events such as the COVID-19 pandemic, and by remote operation. The present work reviews literature that reported the adoption and utilisation of LHD available in the market and presents examples of their practical use. Applications demonstrate the critical role of automation in research development and its ability to reduce human intervention in the experimental workflow. Ultimately, this work will provide guidance to academic researchers to determine which LHD can fulfil their needs and how to exploit their use in both conventional and non-conventional applications. Furthermore, the breadth of applications and the scarcity of academic institutions involved in research and development that utilise these devices highlights an important area of opportunity for shift in technology to maximize research outcomes.

## 1. Introduction

The execution of an experimental task relies heavily on human work. A specific task can become more or less challenging depending on the level of expertise and experience of the person responsible for its completion (from undergraduate students to postdoctoral researchers and principal investigators). Furthermore, experimentation constitutes only a fraction of each individual’s professional responsibilities, rendering the time allocated to laboratory work highly valuable. It might also be necessary to make further adjustments for people with visible/invisible disabilities who require extended time in the laboratory. For other individuals, laboratory-based experimental work may not even be possible at all[1, 2]. Automated and robotic platforms have been developed to assist laboratory operations with the aim of providing support and solutions for the challenges stated above as well as of enhancing experimentation. Interestingly, it has been reported that the majority (up to 89%) of research[3] can be distilled into smaller tasks or steps within a protocol such as liquid manipulation, labware movement, sample acquisition, which can be performed by an automated platform[3]. Particularly, liquid manipulations such as dilutions, mixing, inoculations, and sample preparation among others can be performed easily using liquid handling devices (LHD). Moreover, LHD can be linked to analytical equipment, such as HLPC for rapid analyses. LHD have been deployed for some time and can serve as excellent solutions to processes that are performed repeatedly and do not undergo regular modifications[4, 5].

Although liquid handling equipment is very useful, research tasks can easily become incompatible with them, especially when protocols need to change or evolve frequently, as this will imply repeated workflow construction or modification. This can easily be accentuated by the difficult-to-learn device-specific Graphical User Interface where specific conditions for each experiment are detailed. Specifically, in a scientific or academic environment, students and researchers often utilise new protocols that are designed to execute a unique task, and some of these protocols are only used once. This is especially pertinent in research conducted by postgraduate students, who perform a specific experiment for a critical part of their resource-limited studies, but do not require that protocol again[6]. Even in such instances of ‘niche’ experiments, an automated platform can provide an easy and robust solution or superior results by thoroughly exploring the design space for the experiment. Furthermore, inclusion of such platforms can boost scientific progress by expanding the capabilities of researchers. Complex or expansive design of experiments (DoE) work allow the study of a multitude of variables and environmental conditions at the same time[7-9]. An automated platform would allow a researcher to undertake these tasks without compromising their physical abilities and health. Furthermore, it allows wider participation by providing access to experimentation for people with disabilities[10]. Relinquishing of the human-centred tasks to automated or robotic platforms ultimately sets aside valuable time to be dedicated to experiment design, analysis, and interpretation of the results.

In addition to the concepts mentioned above, advantages of using liquid handling equipment at times of routine operation, such equipment may play key role to solve or prevent external circumstances that manifest themselves beyond the control of the laboratory management and operation. These external circumstances may hinder the work to be conducted in a laboratory by a potential shutdown of all operations or via restricted access to facilities. These are usually unexpected and unplanned, such as the COVID-19 pandemic (2020 – 2022) that restricted human contact and, consequently, access to non-essential work conducted in laboratories[11, 12]. In 2020, a study performed a survey to determine the impact of COVID-19 pandemic in laboratories[13]. The results of the survey showed that two of the least desirable consequences of the pandemic were that experimental work needed to be stopped and that researchers and scientists did not have access to the relevant facilities to carry out their work. These observations resonated with researchers across the globe and even after the relaxation of the isolation and remote working regulations, some places had to work adopting shift patterns to comply with the distancing rules, which rendered certain types of experimental work impossible to conduct. Reflecting on the struggles experienced during those limitations, it can be agreed that appropriately implemented automated platforms could have prevented complete cessation of operations by compressing the time spent in the laboratory only to the preparation of materials and of the platforms to be utilised, while the actual work to be completed would no longer require the presence of a researcher, thus minimizing the human-human interaction to comply with the regulations for the prevention of COVID-19 infection as well as minimising the human-experiment interaction. To expand on this notion, technical staff placed in strategic locations could have facilitated labware manipulation and populate the liquid handling device’s worktables for ease of operation, similar to the operation of a biofoundry[14]. Consideration of all these factors including but not limited to the varying levels of expertise amongst laboratory users, the necessity for inclusive laboratory practices to support people with disabilities, the prospect of expanding the scope of experiments for all researchers, and the prevention of entire operational shutdowns, renders automated platforms essential to transform any laboratory into a robust, resilient, inclusive, and efficient research facility.

Even though LHD can provide a large number of advantages, LHD also have numerous entry barriers. It is well known that they require to undergo proper maintenance to ensure accuracy and precision. Also, critically different to manual pipettes, LHD require a larger footprint that will be permanently occupied by the device. Most of the time and devices require specialized labware (disposable tips) are required for proper operation[4]. Lastly, the cost of acquisition can be prohibitively. Therefore, it is critical to know which devices are available in the market and how can they be used (and are currently being used) to perform cutting edge research.

The objective of the present work it to perform a literature review on the current liquid handling technologies that are commercially available and on how are they being used in research. Critically, it highlights and reports specific applications of automation employed in biosciences research and how it can reduce the human-experiment interaction to favour the human-design-analysis elements of research. We explicitly use the term bio-based laboratories here to emphasise that we conducted a more general search relating to any applications that utilise biological material rather than restricting the scope specifically to biotechnological uses as we believe all bio-based advances can benefit from the adoption of LHDs and related automation technologies. We expect this work to serve as a guide for future widespread application of liquid handling devices in increasing numbers of academic institutions.

## 2. Commercially Available Automated Liquid Handling Platforms

Laboratory equipment, facilitating all experimental work, can range from a simple weighing scale to a complex flow cytometer. This also applies to liquid handling devices; they, too, range from simple to complex machines. All of them can be categorized as robotic equipment since they are able to perform tasks on their own in an independent pre-programmed fashion, but their level of automation is different. The level of automation relates to the amount of tasks a single piece of equipment can perform, to what happens with the information it records (data and metadata), and the propensity of the platform towards flexibility to change. In order to set up the scope of this work, equipment to be considered here needs to fit between automation levels 5 and 6 for liquid handling as described in literature before[4]. These levels comprise equipment that can perform multiple tasks on their own and are flexible enough to allow reconfiguration of protocols and their worktables. Another automation categorization, that includes levels 5 and 6 mentioned before, is the sophistication level ranging from 1 to 4 [15] (with 1 being the highest sophistication level), this relates to the level of decrease of manual inputs. Tier 4 considers devices that only assist pipetting, tier 3 is usually composed of DIY/open-source devices, tier 2 are devices developed to assist specific applications but with enough flexibility to perform other tasks, and tier 1 is for devices than are highly flexible and can provide the most away-time from the experimental work.

Following a detailed assessment of devices available from different commercial suppliers, a summary of the most frequently used existing platforms offered by these suppliers is presented in **Table 1**. This table provides a non-exhaustive compilation of the current main solution providers of liquid handling devices and their most relevant characteristics for the purposes of the present discussion. The devices described here were classified into 3 categories: state-of-the-art, multi-device platforms, and modular equipment. Explained in detail next:

**Table 1.** Collection of commercially available automated liquid handling platforms that can perform liquid handling operations and are sufficiently flexible for different research applications

|  | Brand | Device | Characteristics | Pipette Head(s) | Presence of Robotic Arm |
| --- | --- | --- | --- | --- | --- |
| State-of-the-Art | Tecan | Freedom EVO®, Fluent® | Freedom EVO®: platform tested for liquid handling, and potential multiple liquid handling and gripper arms<br>Fluent®: improves on Freedom EVO® characteristics by allowing an easy workflow construction, flexible and expanded worktable<br>EVOware software: tool for workflow construction and predictive modelling of the expected robotic work | Customizable number of heads between 1 and 384 channels. Available as disposable tips and fixed tips | Yes (Both devices) |
|  | Hamilton | Microlab (Prep®), | Microlab systems: similar in capability, but they | Customizable number of | Yes (Prep® has an add- |
|  |  | NIMBUS®,<br>STAR®,<br>VANTAGE®) | differentiate in the equipment size and module availability. They are capable of identifying labware via embedded cameras to speed up workflow construction<br>The Microlab VANTAGE: complete worktable with multiple arms and modules, can expand into other devices to extend storage and incorporate a secondary deck below the primary one | heads between 1 and 384 channels. Available as disposable tips and fixed tips | on that can connect to the liquid handling arm for plate movement) |
| **Multidevice** | Agilent | Bravo<br>BenchCel<br>Workstation | Workstation composed of Bravo: device for liquid handling; contains a worktable of 9 plate positions<br>BenchCel: device responsible for plate movement and storage (2, 4, or 6 racks of up to 60 plates each). Plate sealer or bar code device can be incorporated | 1 head with option for it between 96 and 384 channels with a volume range of 300 nL to 250 µL | Yes (BenchCel - separate device) |
|  | Analytik Jena | CyBio (Felix, Carry, and QuadStack) | CyBio Felix: liquid handling with a default 9 plate layout, which can be expanded to 15 with a second level worktable. It also allows modules to be attached for specific applications (shaker, thermoshaker, heater/cooler)<br>CyBio Carry: external plate gripper that to transport plate to different plate locations and devices from 80 cm up to 2 m distance<br>QuadStack: storage solution without compromising the deck space Felix composed of 4 rotating stacks of plates with 2 possible heights (555 mm and 755 mm) | 1 head with 1 to 384 channels. 3 types of heads available: An 8 or 12 channel head with a range of 1 to 50 µL, a 1, 16, or 24 channel for a range of 1 to 50 µL, and a 1, 8, or 12 channel for a range of 10 to 1,000 µL | Yes (CyBio Carry - separate device) |
| **Modular** | Gilson | PIPETMAX® | A device with a worktable of 9 positions and only with liquid dispensing capabilities with some attachable modules (orbital shaker, freezer block, and tube racks) | 1 head with 1 to 8 channels for a volume range of 1 to 1200 µL | No |
|  | Bio Molecular Systems | Myra | A small device (< 2 ft<sup>2</sup> and < 10 kg) that provides pressure-based liquid dispensing at low error rates (2% for dispensing 2 µL) and an integrated camera. No capability to move plate positions. | 1 head with single tip head (1 to 50 µL) | No |
|  | Opentrons | OT-2 | An open-source device for high customization. Current modules include shaker, magnetic separation, heater/cooling blocks and a full thermocycler. Contains a worktable with 11 slots for labware or modules | Up to 2 liquid handling arms. Single (1 to 1,000 µL) or 8-channels (1 to 300 µL) | No |
|  | Eppendorf | epMotion series | Simple liquid handler with no labware movement. Application-oriented device configured to performed next-generation sequencing, genomic library preparation, or simple liquid handling. Modules attached depend on the application for each model, but these comprise, vacuum chamber, magnetic separator, and thermal mixer. | 1 liquid arm with 4 volume options (0.2 – 10 µL, 1 – 50 µL, 20 – 300 µL, and 40 – 1,000 µL), either as a single or 8 channels | Some (depends on the model) |
| **Other Devices** | Beckman Coulter | i-Series (Biomek i7) | Single device with all capabilities for liquid handling, but no modules or add-ons available. May come with multiple liquid handling arms | 96 tip head (1 to 300 µL or 5 to 1,200 µL), 384 tip head (0.5 to 60 µL), and the Span-8 pod (8 independent tips and gripper in the same arm) | Yes (Part of the Span-8 arm) |
|  | Perkin Elmer | Sciclone<sup>®</sup> G3 and Zephyr<sup>®</sup> G3 | Sciclone<sup>®</sup> G3: customizable liquid handler that allows only liquids transfer. Zephyr<sup>®</sup> G3: application-specific device for next generation sequencing or solid phase extraction | Up to 2 liquid handling arms. Mandatory a 96 or 384 tip head with option to include an extra 8-channel head | No |
|  | INTEGRA Biosciences | ASSIST PLUS | Device that incorporates independent electronic pipettes into a platform to transform manual work into an automated workflow | 1 with up to 16 channels (Removable electronic pipette). VIAFLO<sup>®</sup> and VOYAGER<sup>®</sup> models with volume ranges (0.5 to 12.5 µL, 2 to 50 µL, 5 to 125 µL, 10 to 300 µL, and 50 to 1250 µL). The D-ONE<sup>®</sup> range from 0.5 to 300 µL or 5 to 1250 µL | No |
|  | Dispensix and Cytena (Both part of BICO) | iDOT and iDOT mini | iDOT: uses a specialised source plate (small apertures in the bottom of the wells) and air pressure to dispense into a destination plate. iDOT mini:same principle but with a single source reservoir | iDOT: 8 liquid channels; source plate up to 96 wells, destination plate up to 1,536 wells; Dispensing is from 8 nL to 50 nL. iDOT mini: 1 liquid channel; single source reservoir with up to 500 µL capacity, same destination plates as iDOT; dispensing is from 8 nL to 500 µL. | No |

### State-of-the-art devices

these comprise equipment that can perform multiple tasks within the same device without further modifications or can integrate additional equipment into the existing space such as liquid handling, labware movement, labware storage, protein/DNA purification, heating/cooling capabilities, and sample analysis among others.

### Multi-device platforms

these are comprised of devices that can perform a single task but can be unified with other single-purpose devices to perform more complicated workflows.

### Modular devices

these are mostly focused in performing liquid movements, but they can incorporate small modules, these are pieces of equipment that can be added to the main worktable such as magnetic separators, heating/cooling devices or thermocyclers to introduce additional capabilities.

Having examined the capability of various platforms, Tecan Group Ltd. and Hamilton Company appear to be the leading suppliers in the state-of-the-art category. Tecan is recognized for its equipment, which can perform liquid handling tasks (from 1 to 96 simultaneous channels) as well as move and store labware. The manufacturers also reported the availability of a range of modular attachments including magnetic separation, vacuum filtration and incorporation of high-throughput chromatography among others to widen the range of tasks that could be executed. An upgrade on this type of device was introduced to provide a more reliable performance and ease of use with updated workflow construction, while promoting integration of third-party devices to design a process without interruptions. Similarly, the range of devices offered by Hamilton provides flexibility on the add-ons that can be attached to their worktables. Both manufacturers provide their proprietary software along with these platforms, which can be used to construct protocols in their native programming languages, but also are able to provide drag-and-drop instructions for a more intuitive construction. Moreover, they can provide 3D simulations of the work the robot is expected to perform.

Another approach adopted by some manufacturers is to reduce equipment cost by providing a multi-device platform. This can be seen from Agilent Technologies, Inc. and Analytik Jena GmbH. These platforms are comprised of multiple devices that are able to run independently, but at the same time are able to integrate seamlessly with one another. In the case of Agilent, this platform is comprised of a device that could work as a standalone liquid handler, another device that is responsible for the movement of plates into a large storage space, which could be enhanced further by a plate sealer and a bar coding device. The ability to provide separate functions and options outside of a single worktable allows the liquid handler to have more space for liquid operations, with large storage in an upright position providing a small footprint for the complete workstation. Analytik Jena has developed a very similar approach by combining devices that work as an independent liquid handling, a carrier and a plate storage; users can opt to acquire only a single robotic platform to speed up experimentation. A complete set, however, can provide more “away time” from the laboratory.

The third frequently encountered approach is to further reduce costs by implementing a single equipment with limitations, which could be enhanced with add-on modules that can be attached to the main worktable. This approach has been adopted by Gilson Incorporated, Bio Molecular Systems, Opentrons and Eppendorf. Gilson and Bio Molecular Systems offer solutions for liquid handling to expedite experimentation with the inclusion of some add-ons. Eppendorf provides small platforms that are fitted for specific applications such as next generation sequencing, DNA library preparation, or for simple liquid handling. To replace the provision of independent modules, suppliers provide packages of the main device with application-specific modules to expedite the required time from acquisition to deployment. Meanwhile, Opentrons offers an open-source solution to the problem with a robot for liquid handling operations, and the add-ons can provide a sterile workplace, recovery of DNA from magnetic beads, a heating/cooling plate, and a complete device to perform a PCR reaction. These add-ons are similar to those typically found in state-of-the-art devices; this means that with less resources it can potentially allow wider access to automation for more users, but it might come with the disadvantage of lacking an intuitive and powerful software as Tecan and Hamilton can provide.

Further devices, which can perform a limited set of instructions, are also available. Beckman Coulter Inc. offers a complete liquid handling platform with no add-ons. PerkinElmer Inc. offers solutions for only transferring liquid between and across 96-well and 384-well plates, and for product purification with scaled-down chromatography columns. INTEGRA Biosciences AG employs a standalone electronic pipette attached to a workbench that can move and perform liquid handling tasks. One final category of devices can only transfer liquids between a single source and a single destination plate (Dispendix GmbH and Cytena). These devices use air pressure or acoustics directed into a well to manipulate very small amount of liquids in the nanolitre scale without pipettes or tips. This makes transfers more accurate and less prone to contamination than its counterparts, which handle liquids via pipetting, but its applications are limited to liquid transfers and require a proprietary source plate with a suitable opening at the bottom of each well.

It is important to highlight other devices that can perform liquid handling are available commercially but are not included here. Such devices as the one provided by Singer Instrument Inc. and Qiagen. These devices perform exceptional tasks, such as colony picking and plating, DNA, RNA, and protein purification, but are not flexible to perform something outside their initial configuration.

## 3. Research Applications Utilising Automated Liquid Handling Devices

This section first presents an investigation of how LHD are used in different areas of application in the literature by implementing a systematic keyword search criterion (Section 3.1) and then moves on to provide an in-depth investigation of the current literature to understand how each LHD was used and how it helped to boost scientific progress (Section 3.2).

### 3.1. Use of LHD in Literature

All liquid handling platforms, regardless of their make, level of automation (tier 5 and 6)[4] or their sophistication tier (1 to 4)[15], have been used to accelerate research and widen the exploration space. Moreover, the complexity of experiments can also be increased through their utilisation. We performed a review of the literature to identify those applications that utilise automated liquid handling devices in bioscience research. For this purpose, we’ve specifically searched for the broad keyword of ‘automation’ along with ‘automated’ liquid handling’ in PubMed (accessed on July 4^th^, 2022). This comparison allowed us to position the prominence of these platforms within the general scope of automation or its lack thereof.

The term “Automation”, defining any general process that can operate by itself minimizing human interaction, was mentioned in 258,030 articles since the time the term was first coined in 1947. The number of publications mentioning the term exponentially increased, particularly since early noughties. (Figure 1a). In contrast, the more specific term “Automated Liquid Handling”, which relates more specifically to the utilisation of automation in biosciences, yielded a search output of only 1,465 articles in PubMed (Figure 1b), of which 285 mentioned uses of one of the commercial platforms presented in Table 1. It is important to consider that “Automated Liquid Handling” does provide a substantial number of articles that utilize a device that is able to move liquids between locations to some extent, but there are also some very niche applications where *de novo* construction of equipment was demonstrated, and the equipment was employed to conduct specific tasks. Such examples include the construction of open-source robots for liquid movement for microscopy applications[16], 3D printed robots for liquid manipulation and controlled using the Internet of Things[17, 18], or the application of liquid handling in microdevices[19, 20], as well as research that was not directly related to bioscience laboratories. Nevertheless, even this initial comparison based on the literature survey that was conducted, was sufficient to demonstrate that the utilisation of automation in biosciences is still at its infancy, with a huge potential as more research laboratories realise the evident advantages of working with such platforms.

**Figure 1.**
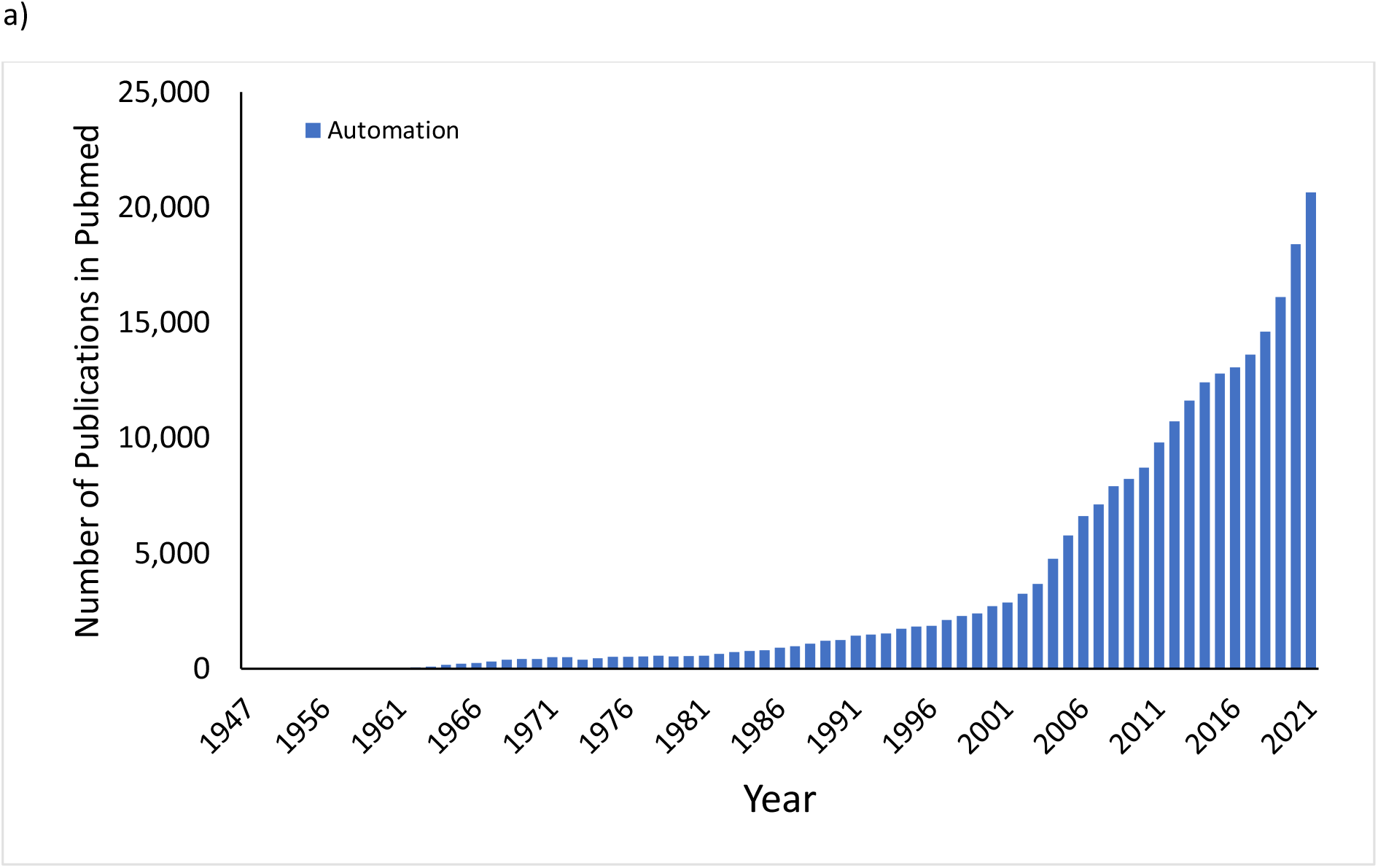
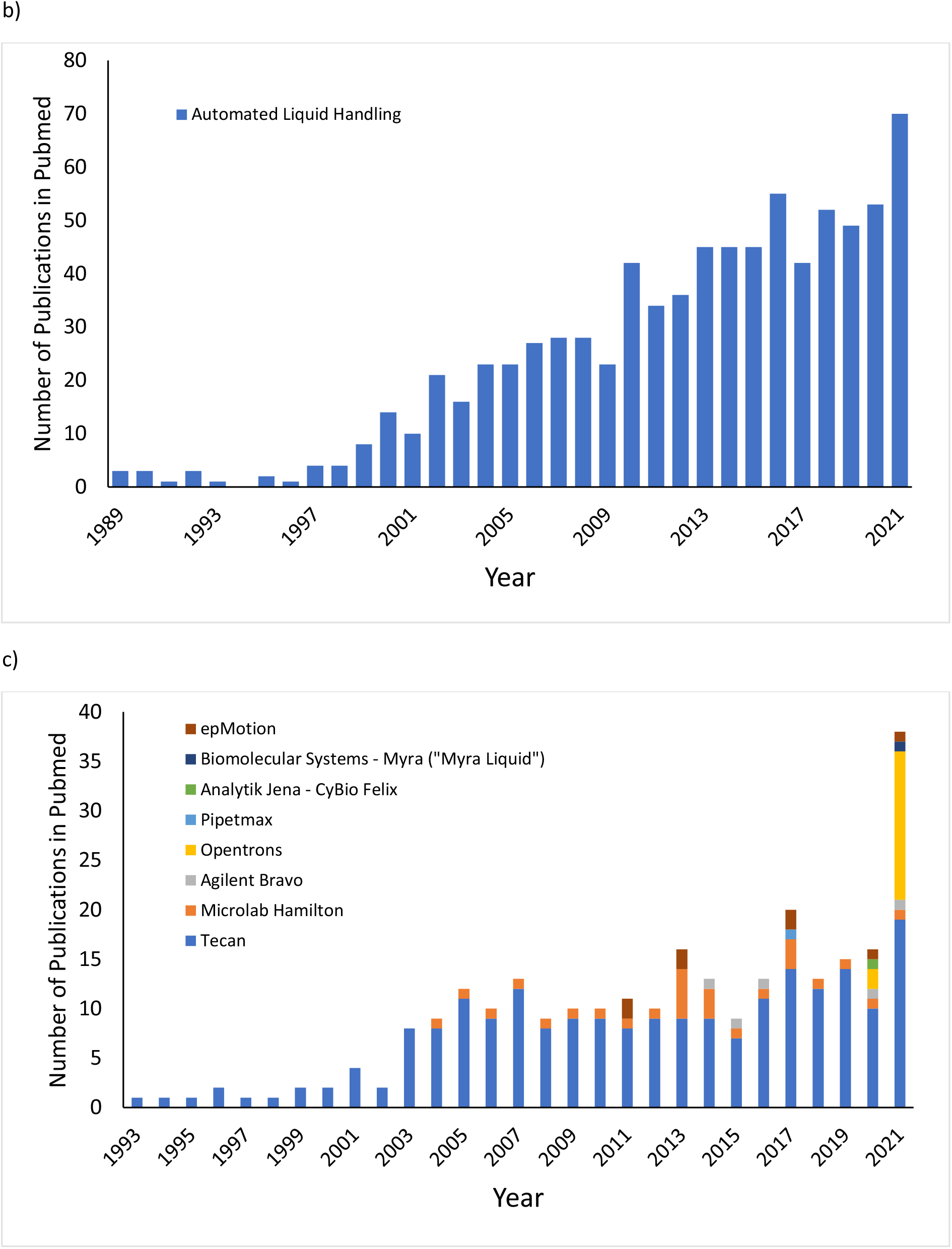
Results for the search in Pubmed using different keywords between the years 1947 (first hit for “Automation”) and 2022. a) shows only “Automation” results, while b) shows results for the specific “automated liquid handling” term. c) shows results for each specific commercial liquid handling device available included in this work.

In order to determine the true use of automated liquid platforms that are of immediate interest, the publications that mention different platforms highlighted in Table 1 were classified accordingly (Figure 1c). This breakdown indicated that the majority of research that utilised automated liquid handling platforms has been conducted using a Tecan device, with the first reports dating back as early as 1993, followed by the Hamilton Microlab devices. A likely reason for this could be that these two platforms were the earliest technologies that were introduced to the market and have evolved substantially over time to improve their performance. A promising development in recent years was the observation that current research also favoured open-source devices as the Opentrons’ OT-2, with 15 articles published in 2021, almost similar to those publications that utilised Tecan platforms in the same year (19 published articles). The remaining platforms have only been sporadically referred to in only one or two publications per year.

### 3.2. Specific Applications of LHD

One of the aims of this work is to highlight how these devices have been used to automate operations and tasks in bioscience laboratories and how they are facilitating research. For this reason, the research articles from the past 3 years (2020, 2021, and 2022 to date) were investigated in detail to demonstrate the types of critical tasks performed while using LHD. This period is specifically selected as it represents the period during which a wider variety of devices (in this case more than three different types) started to appear in the publication landscape, which gives a better representation of the use and the domain of research in which they are used for different devices with a range of technical capabilities and economic burdens. A total of 67 publications were available in the PubMed domain for this 3-year period (2020 – 2022), of which 37 are directly related to an application that utilises an automated liquid handling device (**Table 2)**. Table 2 summarises the information on the device employed, the overall goal of each article, the application area for the liquid handling device, and the institution using it (academia, industry, or a research institution).

**Table 2.** Applications using commercially available liquid handling devices reported in PubMed between 2020 and 2022

| Brand | Device employed | Purpose of research | Use of Device | Type of institution | Reference |
| --- | --- | --- | --- | --- | --- |
| Tecan | Freedom EVO | Develop an automated tube labeller for high-throughput purification | To manipulate the tubes that will be labelled using a Pick-and-Place (PnP) arm | Industry: AbbVie | [31] |
|  | Tecan Fluent and Perkin Elmer JANUS | Perform a contrast between both liquid handling devices to perform toxicological assays | Contrast between 2 devices highlighted the importance of consistent results, but also device bias | Industry: Arup Laboratories;<br>Academia: University of Utah | [53] |
|  | Freedom EVO | Perform a high-throughput process to test purification of viral antigens | Use of the device to expand the research space (samples and variables) applied to scale-down chromatography | Academia: UCL | [21] |
|  | Tecan Fluent and Perkin Elmer JANUS | Identify difficulties and optimization of chromatographic purification in liquid handling devices | To perform scale-down chromatography to identify the bottlenecks and obstacles when trying to set up a new device | Industry: CSL Behring GmbH | [34] |
|  | Fluent | Automate the cultivation of hiPSC and their differentiation into patient-specific retinal pigment epithelial cells for use in drug testing | Functioned as the core of the automated protocol to improve reproducibility | Academia: University of Minnesota | [54] |
|  | Freedom Evo | Demonstrate scheduling complications and solutions for an automated laboratory | Device for sample preparation for deep sequencing | Academia: Nara Institute of Science and Technology and University of Tsukuba; Industry: Tecan Japan Co., Ltd.; Research institution: RIKEN Center for | [37] |
|  |  |  |  | Biosystems<br>Dynamics Research |  |
|  | Freedom<br>EVO | Develop an automated cPass surrogate virus neutralisation test to determine surveillance of SARS-CoV-2 and blocking antibodies | To automate both objectives in order to develop a proof of concept for large-scale application | Industry: Genscript USA, INC., Cayman Chemical, Cure-Hub, FourthWall Testin LCC, DYNEX Technologies, Kaiser Permanente | [22] |
|  | Freedom<br>EVO | Provide with a non-affinity purification method for SARS-CoV-2 spike ectodomain protein | Used along with robocolumns to test multiple operation conditions without wasting time and larger resources | Government:<br>National Institutes of Health | [23] |
|  | Fluent | Set-up a fast and massive Covid-19 testing laboratory | To process all the sample and prepare them for RT-PCR (up to 14,000 per day) | Industry: InActive Blue, Biogazelle, Tecan Trading AG, Sanquin Diagnostics, Bodegro B.V., HiFiBio Therapeutics, Genmab B.V.;<br>Research Institution: Oncode Institute, Hubrecht Institute, University Medical Centre Utrecht, Stichting PAMM, | [24] |
|  | Freedom<br>EVO | Determine antibody levels before and after inoculation with a pneumococcal vaccine (23vPPV) | To validate results obtained from another institution using another automated laboratory | Academia: The University of Melbourne;<br>Research Institutions: Royal Children's Hospital (Melbourne), Population Allergy Research Group & Melbourne Children's Trial Centre | [55] |
|  | Freedom<br>EVO | Develop an automated platform for the quantification of | To perform solid phase extraction using resin-based cartridges and | Academia:<br>Université Paris-Saclay | [33] |
|  |  | succinic acid levels | improve precision percentage |  |  |
|  | Freedom EVO | Validate an automated protocol for quantification of methylmalonic acid in clinical samples | To facilitate a multi-step protocol for sample preparation before using LC-MS/MS analysis | Research Institution: Drug Abuse Laboratory (Sweden) | [36] |
| Hamilton | Microlab ADAP STAR (application-oriented variation) | Create an automated SARS-CoV-2 antibody detection by agglutination-PCR (ADAP) to | To perform complete protocol from sample preparation to qPCR | Industry: Hamilton Company, Enable Bioscience; Research Institutions: Nevada State Public Health Laboratory | [40] |
|  | Microlab STAR | Create an automated islet autoantibody detection by agglutination-PCR (ADAP) for type 1 diabetes | To perform complete protocol from sample preparation to qPCR | Academia: Stanford University; Industry: Hamilton Company, Enable Bioscience | [56] |
|  | Microlab Genomic STARlet (application-oriented variation) | Perform concentration of forensic DNA samples | To incorporate monitored multi-flow, positive pressure, evaporative extraction to perform DNA concentration | Government: Centre of Forensic Sciences (Ministry of the Solicitor General, Canada) | [45] |
| Agilent | Bravo | Automate preparation of libraries for spatial transcriptomic analysis | To provide High accuracy, virtually non-existent technical variability automation for more samples and less processing times than other devices can offer: Complex protocol comprising several steps | Academia: KTH Royal Institute of Technology, Karolinska Institutet | [25] |
|  | Bravo NGS (application-oriented variation) | Perform an automated preparation of single-strand DNA libraries for sequencing ancient biological remains and other sources containing degraded DNA | To manipulate up to 96 simultaneous sensitive samples prone to degradation with minimal cross-contamination | Academia: Max Planck Institute for Evolutionary Anthropology | [44] |
|  | AssayMAP Bravo (application-oriented variation) | Enrich phosphorylated prokaryotic proteins | To capture phosphorylated proteins using microchromatography and concentrate them | Research Institution: Max Planck Unit for the Science of Pathogens | [35] |
|  | AssayMAP Bravo (application-oriented variation) | Develop a high-throughput test for identification of seminal fluid | To perform affinity purification of the semenogelin-II peptide (use of low volumes - 0.0001 uL of semen recovered from casework-type samples) | Academia: Thomas Jefferson University, Arcadia University, The university of Denver, John Jay College of Criminal Justice; Research Institutions: NMS Labs, The Center for Forensic Science Research & Education | [46] |
|  | Bravo | Adapt an automated protocol to accelerate preparation of spatial transcriptomic samples | To facilitate library preparation with 80% reduction in hands-on time for increased number of processed samples | Academia: KTH Royal Institute of Technology | [26] |
| Opentrons | OT-2 | Develop a low-cost and automated platform for DNA assembly (DNA-BOT) | To combine a Biopart Assembly Standard for Idempotent Cloning (BASIC) with a low-cost device (OT-2) to open automation to more users in a biofoundry via accessible configuration | Academia: Imperial College London; Research Institution: London Biofoundry | [57] |
|  | OT-1 | Adapt an open-source robot to manipulate nanolitre volumes | To allow manipulation of nanolitre-scale volumes to reduce reagents consumption | Academia: Brigham Young University; Government: Pacific Northwest National Laboratory | [47] |
|  | OT-2 (contrast with MagMAX<sup>TM</sup>) | Evaluate low-cost protocols for RNA extraction for SARS-CoV-2 detection | To perform in-house RNA extraction for subsequent qPCR and to compare performance with | Research Institution: Hospital Universitario La Paz | [41] |
|  | Express-96 system) |  | outsourcing for resource utilisation |  |  |
|  | OT-2 | Optimise large-scale diagnostics for SARS-CoV-2 using RT-PCR | To process large number of SARS-CoV-2 detection test using 5 testing lines, with 2 OT-2 each to test up to 5,460 samples per day, selected for low-cost and flexibility of programming | Research Institution: Hospital Universitario Virgen del Rocío | [27] |
|  | OT-2 | Create a semiautomated preparation for isobaric tag samples | To automate approximately 60% of the complete protocol facilitating liquid mixing, magnetic separation, and miniaturized chromatography | Academia: Harvard Medical School | [38] |
|  | OT-2 | Develop an automated platform for SARS-CoV-2 testing using RT-qPCR | To use an open-source device to configure (changes to liquid classes, aspiration heights, creation of a log system, and custom labware) and adapt to a clinical setting | Academia: University of Barcelona; Research Institution: Hospital Clínic de Barcelona | [42] |
|  | OT-2 | Create a complete open-source modular framework applied to automated liquid handling and imaging | To synchronize with a custom-made microscopy system fitted into the device and collect data in a repository | Academia: KTH - Royal Institute of Technology, University of Bath, Friedrich-Schiller-Universität Jena, Uppsala University; Research Institution: Leibniz Institute for Photonic Technology | [48] |
|  | OT-2 | Automate shotgun proteomics | To demonstrate the use of a low-cost platform for preparation of proteomic samples and easy-to-share instructions to other OT-2 devices | Academia: University of Colorado | [39] |
|  | OT-2 | Adapt an existing workflow to manipulate large | To demonstrate the applicability of a previous protocol | Research Institution: Hospital | [28] |
|  |  | volumes for SARS-CoV-2 RT-PCR | design and confirm prediction results | Universitario Virgen del Rocío |  |
| Gilson | Pipetmax | Automate Darwin Assembly using Synthace | To accelerate workflow construction, while reducing errors and hands-on time | Academia: KU Leuven, University College London; Industry: Synthace Ltd, Sangamo Therapeutics Inc.; Research Institution: Medical Research Council Laboratory of Molecular Biology | [50] |
| Analytik Jena | CyBio-Well Vario 96 (application-oriented variation for 96 simultaneous channels) | Identify proliferation factors in MCF7 breast cancer cells using RNA interference | To dispense siRNA into experimental plates | Academia: Georgetown University, Drexel University, Kazan Federal University; Research Institution: Fox Chase Cancer Center | [58] |
|  | CyBio Felix | Perform rapid design and implementation of a SARS-CoV-2 test in an NHS diagnostic laboratory | To perform RNA extraction from clinical samples in conjunction with other platforms | Academia: Imperial College London; Industry: Riffyn Inc.; Research Institution: London Biofoundry, North West London Pathology, Charing Cross Hospital, Hammersmith Hospital Trust | [29] |
| | CyBio Felix | Develop and validate immunoassay to detect APP669-711(Aβ-3-40) | To perform the immunoprecipitation using magnetic beads | Academia: Georg-August-University, Friedrich-Alexander-University of Erlangen-Nuremberg, Technische Universität Dresden, University of Aveiro; Industry: Tecan Group Company, Roboscreen GmbH; | [59] |
|  |  |  |  | Research<br>Institution: Max-Planck-Institute of<br>Experimental<br>Medicine, German<br>Center for<br>Neurodegenerative<br>Diseases |  |
| Biomolecular<br>Systems | Myra | Create a mobile<br>SARS-CoV-2<br>laboratory for in-<br>field testing | To allow portability<br>and high-throughput<br>processing | Academia: the<br>University of<br>Western Australia;<br>Research<br>Institution:<br>PathWest<br>Laboratory<br>Medicine WA,<br>National Health<br>Laboratory | [43] |
| Eppendorf | epMotion | Construct a<br>foundation to<br>perform routine<br>testing for SARS-CoV-2 detection | To perform RNA<br>extraction of clinical<br>samples | Research<br>Institution:<br>Auckland City<br>Hospital | [30] |

Each article had a particular objective for the research carried out and the liquid handling device corresponds to a tool employed as a key component for the completion of the work, or a part of it. Moreover, the inclusion and utilisation of a liquid handling device was decided on the basis that it helped to solve a specific problem or overcome certain challenges. Most of the applications shown in Table 2 are related to the ability to process a large number of samples in a short period of time[21-31]. This is one of the most evident technical advantages of a liquid handling device to be used for the execution of a simple task, i.e., dispensing of some liquid into a specific location, which can become time-consuming when performed at scale. In one example, the testing for SARS-CoV-2 virus was performed for up to 5,460 samples per day[27]. As a hypothetical scenario, assuming it requires a single aspiration-dispense cycle with a hypothetical 1 second for each step plus 1 second for pipette movement for the same number of samples, it will require approximately 4.5 hours to execute this single task if a single-channel pipetting instrument is used. Extrapolation of this simple example to systems, which are able to handle up to 14,000 samples per day, outlines a very clear overview of how critical such systems are as throughput increases[24].

Speeding up the operations, particularly for high number of samples is a key advantage, but it is far from being the sole advantage of using automated liquid handling platforms and sometimes, depending on the experiment carried out, time constrains their use[32]. One clear advantage of these platforms lies in the precision of their technology. The use of multi-channel pipetting with precise flow control has allowed scale-down chromatographic experiments to be performed. Miniaturized columns have not only allowed to accelerate the discovery of adequate purification conditions for pharmaceutical products[21, 33], but also to identify key bottlenecks and obstacles before performing large-scale experimentation[34]. Cartridge-based purification has also been applied for solid-phase extraction[33], enrichment of phosphorylated proteins[35] and quantification of metabolites in clinical samples[36].

Other applications for the liquid handling devices discussed here focussed on the preparation of a large number of samples for subsequent analyses, such as sequencing[37], spatial transcriptomics[25, 26], isobaric tagging (from liquid mixing to miniaturized purification)[38] and shotgun proteomics[39]. Given the nature of analyses applied to samples containing genetic material, the application of a liquid handling devices allows to process a large number of samples to improve the results.

A similar time-critical application for a large number of samples has been the use of these devices for the detection of SARS-CoV-2 virus in clinical samples. In the past 3 years, most devices, but especially small and less expensive ones, had been featured in at least one published application where they have been used to create an automated platform or improve existing ones. Analytik Jena’s CyBio Felix[29] and Eppendorf’s epMotion[30] were used to perform RNA extraction from patient samples in clinical settings. Other devices were employed to perform more complex applications related to SARS-CoV-2 virus than those highlighted above. A Freedom EVO was used to track SARS-CoV-2 infection through the presence of antibodies using a cPASS Surrogate Virus Neutralizing Test (sVNT)[22], while a Hamilton Microlab STAR, in its application-oriented version for Antibody Detection by Agglutination-PCR (ADAP), was used to survey antibody prevalence in samples[40].

The impact of SARS-CoV-2 in our day-to-day operations also helped improve science and paved the way to demonstrate new ways of thinking on how to enhance laboratory capabilities. Opentrons’ OT-2 has been used to demonstrate that in-house RNA extraction and qPCR processing can decrease economic burden to some laboratories to replace the outsourcing of such workflows[41]. In another study, SARS-CoV-2 samples collected from different sources was considered a challenge since the variation in these sources manifested itself as differences in sampling tubes, liquid levels, and liquid viscosity. The open-source nature of the OT-2 was exploited to develop an easy-to-configure device that can change liquid classes, aspiration heights, create a log system and employ custom labware[42]. Yet another example focused on the compromise between capabilities, accessibility, affordability, and footprint for different applications was demonstrated with a Myra device when it was incorporated into a mobile laboratory to perform RNA extraction to replace manual execution of the protocol[43]. This resulted in superior mobility without compromising on the high-throughput processing capabilities.

Automated platforms can also provide help with delicate tasks where human interaction can degrade or contaminate samples. An example of such a challenge is observed in the analysis of ancient biological samples with fragile and heavily degraded DNA that needs to be processed. When challenged by such a sample, the liquid handling device allowed to decrease cross-contamination during the processing of multiple samples simultaneously[44]. The same principle and the requirements apply to samples used for criminal analyses. The use of liquid handling devices allowed to perform complex multi-step protocols to concentrate DNA successfully in a publication by Chase, *et al*. Prior to the utilisation of such devices, if the concentration of the sample was lower than a certain threshold, then the analysis would not be carried forward as the risk of cross-contamination substantially impaired the validity of the results[45]. The increase in the capabilities of the device and to be able to work with low volumes via concentration was also highlighted by Davidovics *et al*., who demonstrated the successful utilisation of low volume (below 0.0001 µL) semen samples to perform purification of the semenogelin-II peptide for criminal identification[46].

Lastly, it is critical to highlight that the devices need to be able to adapt and be reconfigured as per changing user requirements. This has a massive beneficial impact on research and, successful examples of such application adjustments are already available: Opentrons’ OT-1 was modified to manipulate nanolitre-scale volumes (50 nL) with less than 3% error[47] and an OT-2 was fitted with a mobile and custom-made microscope to automate imaging[48]. Before any research application is automated, the impact of the utilisation of a liquid handling device can be analysed by assessing scheduling and task completion[37]. Depending on the task deployed, this analysis is critical as it allows the exploration of different configurations and identify better options for the operation to be executed than those already evaluated. An example of how relevant scheduling becomes was observed in the utilisation of liquid handlers for SARS-CoV-2 testing. With automation, it was expected that all samples arriving at a test facility could have been processed daily provided that the capacity of the liquid handling platforms was not exceeded, but this was not necessarily reported to be the case, with the platforms reporting up to 90% of the results within 24 hours[28]. While this is a fast turnover, provided the restrictions imposed by the pandemic setting, an improved scheduling might be implemented as required depending on the type of the application.

All the applications reviewed here (**Figure 2**) have facilitated research in a specific way and especially open-source devices can make automation more accessible and flexible for non-conventional applications. Still, there is a learning curve in the creation of coherent workflows that can perform the desired tasks in an efficient way. In order to expedite this, dedicated software has been developed to cross communicate between devices and make workflow construction more intuitive[4]. The application of such web-based software such as Synthace has been reported to build a Darwin Assembly (generation of single-stranded DNA template, isothermal assembly, and amplification for cloning[49] in a Gilson Pipetmax[50].

**Figure 2.**
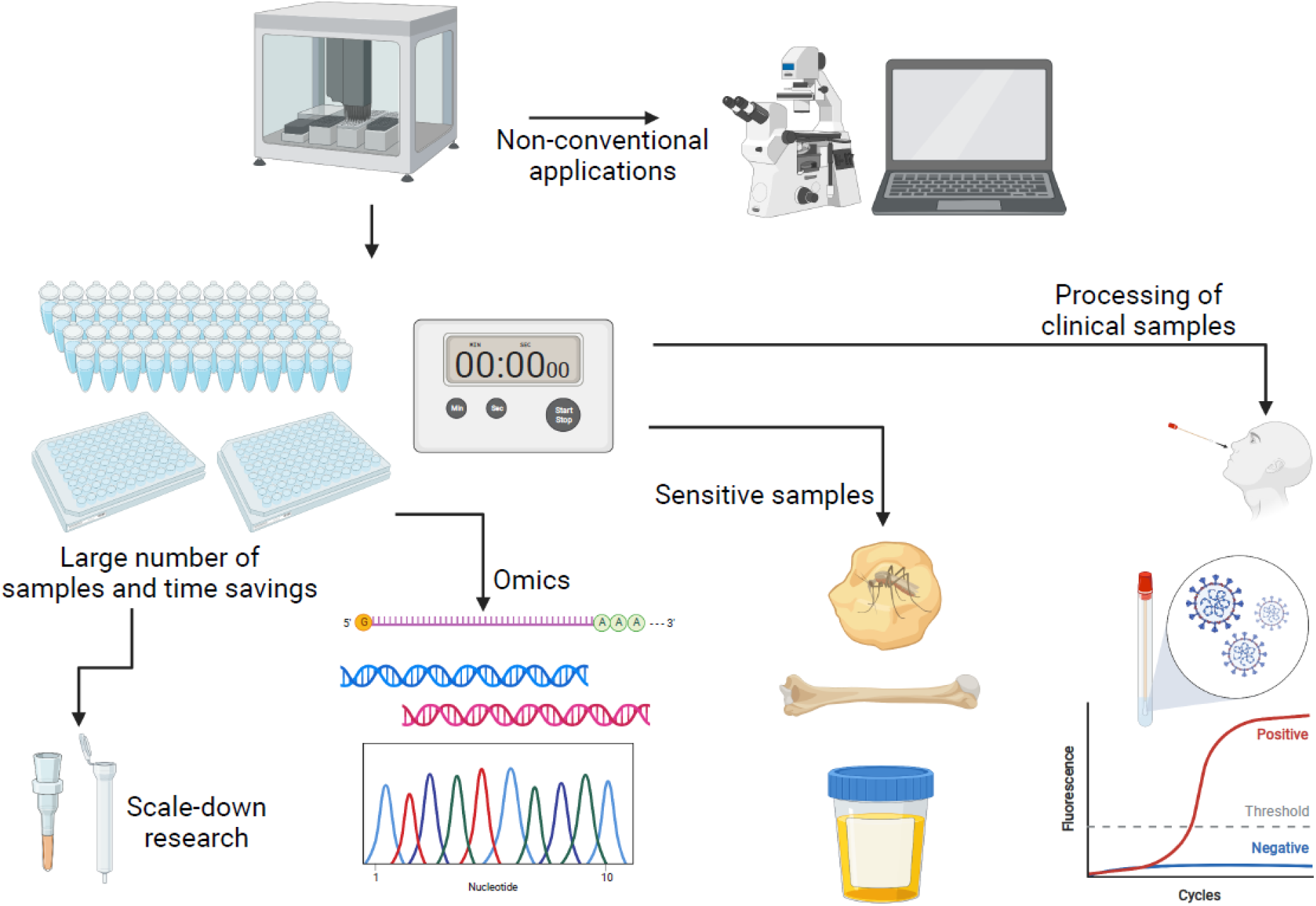
Current applications (2020 – 2022) for liquid handling devices. Common applications include large sample processing, omics analyses, processing of sensitive, delicate, and forensic samples, and expedite clinical samples evaluation (i.e., SARS-CoV-2). Additional non-conventional applications (DIY) include integration of microscopic imaging, nano-litre scale modification, and modelling tools. Created with BioRender.com

It is important to note that only 23% of these automation-enabled research highlighted above were carried out entirely by academic institutions highlighting the resource and access limitations associated with the utilisation of automated liquid handling platforms in a higher education setting. Collaboration with research institutions and industrial partners appears to be key for academic access to this equipment across the world given the capital cost barriers to access. Some of the research discussed above explicitly acknowledged collaborators from industry, who are in charge of these devices in question. From an academic institution’s point of view, it is important to identify key use of these systems, explore their applicability in different settings and how this flexibility could lead to increase in the number of users as well as widening the user application domain in order to justify the investment. Current developments in low-cost and open-source instruments can open automation to more users and institutions to perform ground-breaking research. Budget limitations render it critical for academic institutions to analyse the cost of devices. For this reason, different levels of automation or complexity, which could assist laboratory operations to different extent, are presented here. Operating costs of these devices are also closely related to how energy-efficient they are. The use of LHD typically require extended work time that can increase energy-associated costs. Finally, costs can be aggravated by geographical location of academic institutions. Suppliers of these devices are located typically in USA or Europe as major academic and industry hubs, but other countries need to consider importation fees. All these aspects are critical to the decisions to be made around the selection of an appropriate device.

Within the literature reviewed here, it is important to highlight that LHD are typically used as a tool or device in the laboratory and the analysis of their performance is not presented. LHDs are important tools but they are susceptible to errors caused by wear of the mechanical pieces or by human input. This needs to be assessed before any deployment. Previous literature that analysed such errors showed how variable results could be. Using a contrast between manual and automated work in liquid chromatography coupled with mass spectrometry, UV profiles demonstrated to be quite similar giving the confidence of similar performance[8]. Possibility of errors and the improvement of reproducibility are important factors in every work, but are especially important in non-conventional applications, such as when applying liquid manipulation in point-of-care testing systems[51]. These applications can introduce human error by non-trained users. A recent work developed a camera-based protocol to determine the error introduced by the addition of reagents into the samples provided[52]. Another non-conventional application mentioned previously is the modification of an OT-2 device to manipulate nanolitre volumes[47]. In this work it became critical to demonstrate that error minimization and reproducibility could still be achieved despite using the device outside of the normal operating range.

## 4. Looking ahead

The use of automation and, particularly, automated liquid handling platforms reduce experimental time and errors. Using these platforms, large experiments and big datasets can be completed without the intervention of a person, which allows researchers to focus on planning and data analysis. These devices allow the manipulation of sensitive and critical samples, and those that are prone to degradation in a safe and efficient manner. Moreover, with the proper expertise most devices, especially open-sourced ones, can be easily adapted to perform non-conventional tasks to enhance the research capabilities of their users. LHD can be further utilized and expanded to investigate a diverse range of areas, such as the development of new biomedical devices, reduction of waste during experimentation and subsequently during actual large-scale applications and maximize analysis of green technology developments.

The existence of entry barriers to automation is being reduced everyday with new device developments, the ability to share workflows, and the design of dedicated software that allows cross-communication. COVID-19 pandemic further pushed research into exploring automation-friendly environments to provide everyone with access, even remote access, to a laboratory setting.

Reflecting on the current standing of automated liquid handling systems in experimentation, we are increasingly exploring different ways of incorporating automation into our own research practices. We are approaching this both from the perspective of enhancing experimental research as well as producing graduates, at all levels, with the skills they will need in modern industrial research laboratories.

One particular focus for us is the evaluation of the strengths and weaknesses of different systems and approaches in adopting automation in tailored and customised research applications, along with the routine operations that are carried out on a regular basis. Our efforts primarily focus on the investigation of the entry barriers for researchers to start adopting these technologies and we seek measures and technologies that generate minimal hesitation and resistance across the widest possible range of users of such platforms. Automation of our bioscience research practices not only sit at the core of our efforts in achieving the digital goals of Industry 4.0 initiative in an academic setting, but also supports our transition into an Industry 5.0-compliant future.

## Acknowledgments

The authors would like to acknowledge funding from University College London’s Sustainable Physical and Digital Places for Education and Research (SPiDER) group and support from EPSRC CDT Bioprocess Engineering Leadership (Grant Number EP/L01520X/1). For the purpose of open access, the authors have applied a Creative Commons Attribution (CC BY) licence to any Author Accepted Manuscript version arising. Authors have no conflict of interest to declare.

